# Sexual receptivity increases in synchrony with the ovulatory cycle in female medaka

**DOI:** 10.1101/2024.08.06.606788

**Authors:** Soma Tomihara, Rinko Shimomai, Mikoto Nakajo, Yoshitaka Oka, Chie Umatani

## Abstract

Successful reproduction requires coordinated regulation of gonadal function and sexual behavior. However, the mechanisms of such a regulation remain elusive. Here, we used a teleost medaka to find out possible involvement of ovulation in the control of female sexual behaviors by analyzing the sexual behaviors of the female gene-knockout medaka that do not ovulate and found that ovulation is essential for the facilitation of female sexual receptivity. Behavioral recordings and anatomical examinations showed that the sexual behavior of medaka occurs only after the ovulation. Furthermore, progesterone administration partially reinstated the sexual receptivity of the anovulatory knockout females. Taken together with the result that progesterone receptor is expressed in the brain regions that are considered strong candidate for regulation of sexual behavior, we propose that female sexual receptivity is facilitated in synchrony with the ovulatory cycle via progesterone receptor signaling in specific brain regions that occurs around the timing of ovulation.

## Introduction

In most vertebrates, males and females perform innate sex-specific sexual behaviors when both sexes possess mature gametes in their gonads, which is essential for successful reproduction. Mammals usually show specific ovulatory cyclicity, which has been shown to be regulated by hypothalamic-pituitary-gonadal (HPG) axis and kisspeptin neuronal systems. Hypothalamic kisspeptin neurons regulate both pulsatile and surge release of gonadotropin-releasing hormone (GnRH)^1,2,3,4^. Pulsatile release of GnRH promotes the release of follicle-stimulating hormone (FSH) and luteinizing hormone (LH) from the pituitary, resulting in the early follicular development in the ovary^5,6,7^. On the other hand, GnRH surge triggers LH surge from the pituitary, which leads to the late follicular development and ovulation^8,9^. Accumulating evidence in rodents has also shown that serum concentrations of some sex steroid hormones show regular fluctuations, which generate cyclic patterns of folliculogenesis and ovulation, i.e., the ovulatory cycles^7,10,11^. It has been generally accepted that the ovulatory cycles contribute to the regulation of sexual receptivity of the female, the motivation for mating with the male. On the other hand, the neuroendocrine regulatory mechanisms that generate ovulatory cycles as described above have been mainly studied in mammalian species, which are considered to be biologically different in many aspects from the other non-mammalian vertebrate species^8,12^. Therefore, it is necessary to elucidate whether and how the activation of sexual receptivity is regulated by the ovulatory cycles in non-mammalian vertebrates.

Teleosts are known to comprise the major group of non-mammalian vertebrates with a great biodiversity and have served as nice model animals especially for the study of instinctive behaviors such as sexual behaviors or the endocrine/neuroendocrine regulation of reproduction. There are studies that analyzed the frequency of spawning and estimated the timings of oocyte maturation and ovulation. For example, a teleost bambooleaf wrasse (*Pseudolabrus japonicus*) is known as a daily spawner and have diurnal maturation cycle of oocytes^13^. Another teleost, mummichog (*Fundulus heteroclitus*), is known to spawn daily during its breeding season from late March to August, and the females show daily ovulatory cyclicity^14^. These studies suggested that the ovarian states may affect the activation of female sexual behavior in these teleosts. However, the neuroendocrine mechanisms underlying the synchrony of ovulation and sexual behavior have not been studied in detail, mainly because of the difficulty in applying various molecular genetic tools and contemporary neurobiological techniques to these teleosts. Accordingly, it has been unclear whether and how the activation of sexual receptivity is linked to the ovulatory cycles.

In the present study, we used a teleost medaka (*Oryzias latipes*). Medaka has a 24-hour ovulatory cyclicity, and it is rather easy to analyze the dynamics of ovulation. We can also take advantage of being able to apply molecular genetic techniques^15,16^, and neuroendocrine mechanisms underlying the ovulatory cycles have been analyzed by using medaka. Gene knockout (KO) analysis has demonstrated that FSH-deficient female medaka possess small ovary with immature oocytes because of the disrupted folliculogenesis, and GnRH/LH-deficient females do not ovulate despite normal folliculogenesis, leading to infertility of these females^17^. Such results suggest that FSH and LH contributes to folliculogenesis and ovulation, respectively, although only LH release is under the control of GnRH signals from the GnRH neurons in the preoptic area. It should be noted that accumulating evidence suggests that kisspeptin neuronal systems in non-mammalian vertebrates are not involved in the HPG axis regulation but rather in the control of sexual behaviors^12,18^. Thus, it has recently been postulated that the neuroendocrine systems for regulating ovulatory cycles in teleosts are different from those in mammals. Therefore, it is worth analyzing the relationship between ovulatory cycles and sexual behavior in a model teleost medaka, which will be helpful for the deeper understanding of the general neuroendocrine mechanisms for the activation of sexual receptivity in synchrony with the ovulatory cycles in vertebrates. Another advantage of the use of medaka is that the male and female medaka pairs show stereotypical sexual behavior repertoires, which can be readily analyzed quantitatively^19^. Here, we analyzed the sexual behavior of the female gene-knockout medaka that do not ovulate and found that they do not show sexual receptivity for males. We also analyzed and determined the timeline of ovulation and sexual behavior of medaka pairs before and after dawn. Finally, we obtained behavioral and neuroanatomical results to suggest that female sexual receptivity is facilitated in synchrony with the ovulatory cycle via progesterone receptor signaling in specific brain regions that occurs around the timing of ovulation.

## Results

### Anovulatory gene knockout females do not exhibit clasping

To investigate the relationship between the ovarian state and the female sexual behavior, we used some lines of gene knockout medaka showing abnormal ovarian states (*fshb*^-/-^, *lhb*^-/-^, and *gnrh1*^-/-17^) and analyzed their sexual behaviors. To begin with, we analyzed sexual behavior of *fshb*^-/-^ female medaka, which exhibit immature ovaries, paired with a wild type (WT) male. We analyzed the frequency of following, the percentage of time spent for following, and the frequency of courtship display, all of which are regarded as indices for male motivation of sexual behavior towards females. There were no significant differences in all three indices between *fshb*^+/+^ and *fshb*^-/-^ females (Figures 1A-C), suggesting that the male sexual motivation is not affected by the ovarian maturational states of the female. On the other hand, we found significant differences in the Kaplan-Meier curves indicating the latency to the first clasping/spawning, which indicates the sexual receptivity of the female (Figures 1D and 1E). We also found that *fshb*^-/-^ females never spawned, as expected from their immature ovary. They did not show clasping and appeared to reject to be wrapped by male for clasping. These results suggest that ovarian maturation affects the sexual receptivity of the female medaka, while leaving the male sexual motivation unaffected.

**Figure 1.**
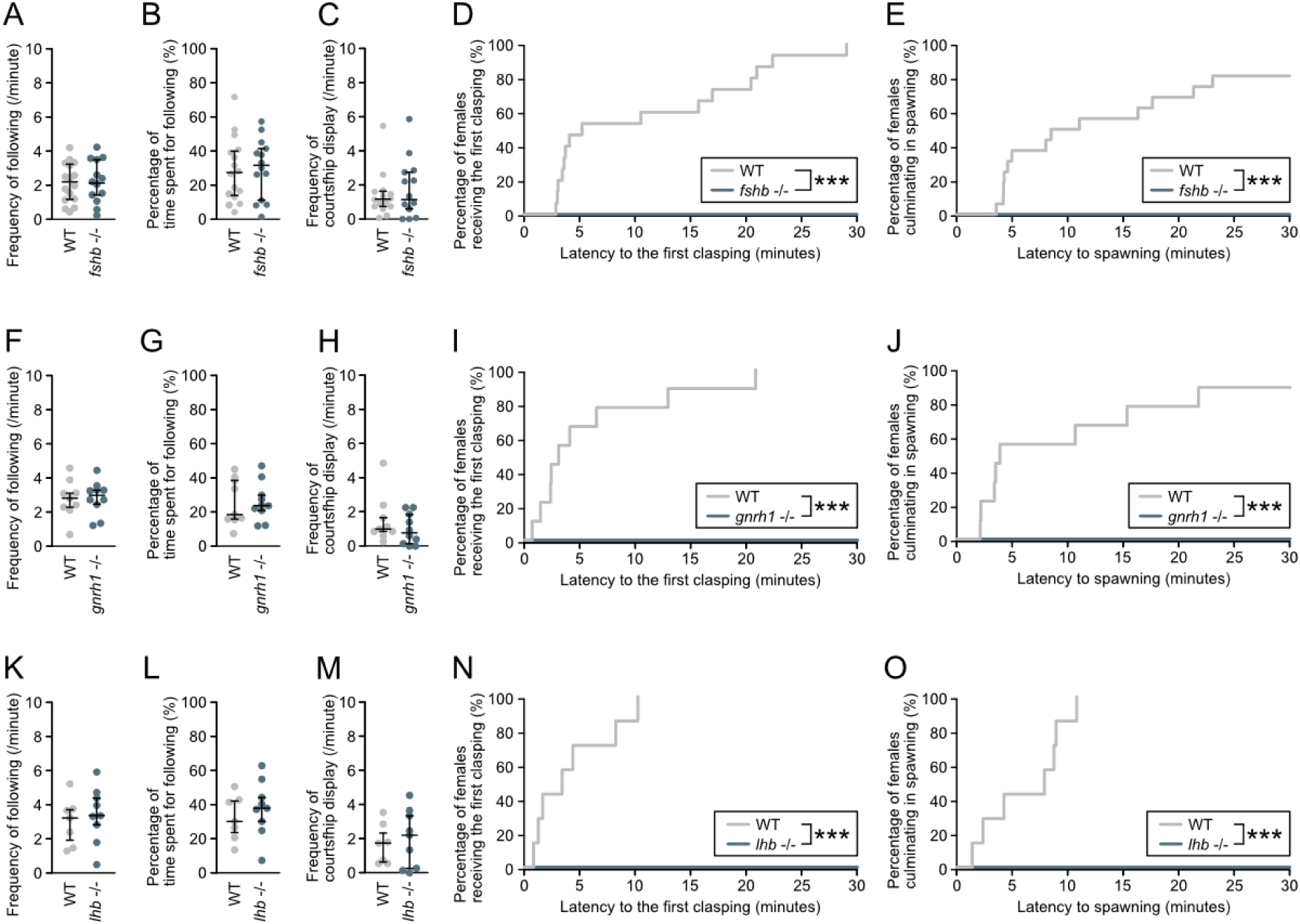
Female KOs of the HPG axis-related genes do not exhibit sexual receptivity. *fshb*^+/+^ (n = 16), *fshb*^-/-^ (n = 14), *gnrh1*^+/+^ (n = 9), *gnrh1*^-/-^ (n = 10), *lhb*^+/+^ (n = 7), and *lhb*^-/-^ (n = 9) females were tested for mating behavior with a wild type (WT) male. *fshb*^+/+^ , *gnrh1*^+/+^ , and *lhb*^+/+^ were indicated as WT. Frequency of male following of *fshb*^-/-^ female (A), Percentage of time spent for male following of *fshb*^-/-^ female (B), Frequency of male courtship display toward *fshb*^-/-^ female (C), reverse Kaplan-Meier plot indicating the percentage of *fshb*^-/-^females receiving the first male clasping (D) (\*\*\**P* = 0.0000002), and reverse Kaplan-Meier plot indicating the percentage of *fshb*^-/-^ females culminating in spawning (E) (\*\*\**P* = 0.000007). Frequency of male following of *gnrh1*^-/-^ female (F), Percentage of time spent for male following of *gnrh1*^-/-^ female (G), Frequency of male courtship display toward *gnrh1*^-/-^ female (H), reverse Kaplan-Meier plot indicating the percentage of *gnrh1*^-/-^ females receiving the first male clasping (I) (\*\*\**P* = 0.000004), and reverse Kaplan-Meier plot indicating the percentage of *gnrh1*^-/-^ females culminating in spawning (J) (\*\*\**P* = 0.00006). Frequency of male following of *lhb*^-/-^ female (K), Percentage of time spent for following of *lhb*^-/-^ female (L), Frequency of male courtship display toward *lhb*^-/-^ female (M), reverse Kaplan-Meier plot indicating the percentage of *lhb*^-/-^ females receiving the first male clasping (N) (\*\*\**P* = 0.00002), and reverse Kaplan-Meier plot indicating the percentage of *lhb*^-/-^ females culminating in spawning (O) (\*\*\**P* = 0.00002). We used Mann-Whitney *U* test for the analyses of frequencies and percentages (A-C, F-H, L-M), and log-rank test for the analyses of difference of reverse Kaplan-Meier plots (D, E, I, J, N, O).

Since *fshb*^-/-^ females show severe deficiency in ovarian maturation comparable to that in non-breeding status, it remains unclear which events in the ovarian status, oocyte maturation, ovulation, etc., caused abnormal behavior in *fshb*^-/-^ female. Therefore, we next focused on *gnrh1*^-/-^ females and *lhb*^-/-^ females, which are infertile because both of them are anovulatory in spite of the fact that they show full follicular growth and normal gonadosomatic index. To investigate the possible relationship between female receptivity and ovulation, we analyzed the sexual behavior of *gnrh1*^-/-^ and *lhb*^-/-^ female paired with a WT male. There were no significant differences in the frequency of following, the percentage of time spent for following, and the frequency of courtship display (Figures 1F-1H), suggesting that males normally approached and courted to *gnrh1*^-/-^ females. However, *gnrh1*^-/-^ females never showed clasping and spawning, whereas almost all *gnrh1*^+/+^ females showed the first clasping within 15 minutes after they started to interact (Figures 1I and 1J). Similarly to *gnrh1*^-/-^ females, no significant differences were found in the frequency of following, the percentage of time spent for following, and the frequency of courtship display of *lhb*^-/-^ females (Figures 1K and 1M), suggesting that *lhb*^-/-^ females were normally courted by males. Nevertheless, *lhb*^-/-^ females did not show either clasping or spawning. There were significant differences between *lhb*^+/+^ and *lhb*^-/-^ females in the Kaplan-Meier curves showing latency to the first clasping/spawning (Figures 1N and 1O).

In order to confirm that these results were caused by the lack of ovulation, we rescued the ovulation by the intraperitoneal administration of pregnant mare serum gonadotropin (PMSG) to *gnrh1*^-/-^ females and analyzed their sexual behavior with WT males (Methods are described in Document S1). Two days after PMSG administration, three of four pairs comprising PMSG-administered female and WT male (PMSG+ pairs) showed clasping, and two pairs normally spawned (Table 1 and Figure S1). The fertilization rate was 46.7% (7/15 eggs) and 97.0% (32/33 eggs), respectively. Furthermore, all of the four PMSG+ pairs performed clasping three days after administration, and two of four PMSG+ pairs spawned within 30 minutes of interaction, and one PMSG+ pair spawned approximately six minutes after the start of analysis (Fig. S1) with fertilization rates of 81.8% (9/11 eggs), 88.9% (24/27 eggs), and 100% (4/4 eggs), respectively. Also, it should be noted that male motivation of courtship to females was not affected regardless of the presence or absence of their ovulation. From these results, it is suggested that ovulation is closely related to the activation of female receptivity.

**Table 1.**

| **Days after**<br>**injection** | **Pairs showing clasping / Total** |  |  | **Pairs culminated in spawning**<br>**/ Total** |  |
| --- | --- | --- | --- | --- | --- |
|  | **+ PMSG** | **+ PBS** |  | **+ PMSG** | **+ PBS** |
| **1** | **0/4** | **0/4** |  | **0/4** | **0/4** |
| **2** | **3/4** | **0/4** |  | **2/4** | **0/4** |
| **3** | **4/4** | **0/3** |  | **2/4** | **0/3** |

### Ovulation occurs about two hours before the lights-on

We next focused on the temporal relationship between ovulation and activation of female sexual receptivity. Since there have been only a few studies in medaka that analyzed the timing of ovulation *in vivo*, we analyzed the temporal changes in the number of spawned eggs and the ovarian morphology to detect the timing of ovulation (Figure 2A). First, we counted the average number of spawned eggs during three days before the behavioral experiment, which was defined as the number of eggs attached to the female abdomen (Figure 2B). Since the previous study assumed that ovulation occurs at around 7:00 AM (1 hour before lights-on)^25,26^, three WT females were dissected every hour from 11:00 PM to 7:00 AM, and their ovaries were morphologically analyzed. Ovulated eggs, which were characterized by the fibrous membranes (arrowheads in Figure 2C), were not found between 11:00 PM and 4:00 AM except for two ovulated eggs from the female that was dissected at 2:00 AM (average number of spawned eggs during the last three days before dissection was eleven) (Figure 2B). On the other hand, ovulated eggs were found in the females that were dissected at 5:00 AM, and each percentage of ovulation ((the number of ovulated eggs / the average number of spawned eggs during the last three days) × 100) for the three females was 10%, 9.1%, and 64%, respectively. In addition, the percentages of ovulation at 6:00 AM and at 7:00 AM were almost > 70% (Figure 2B). Consistent with these results, ovulated eggs, which were located in the ovarian cavity, could be detected from the dorsal side of the ovaries that were dissected out after 5:00 AM (Figure 2C). These results suggest that ovulation occurs between 5-7 AM, approximately two hours before the lights-on (8:00 AM).

**Figure 2.**
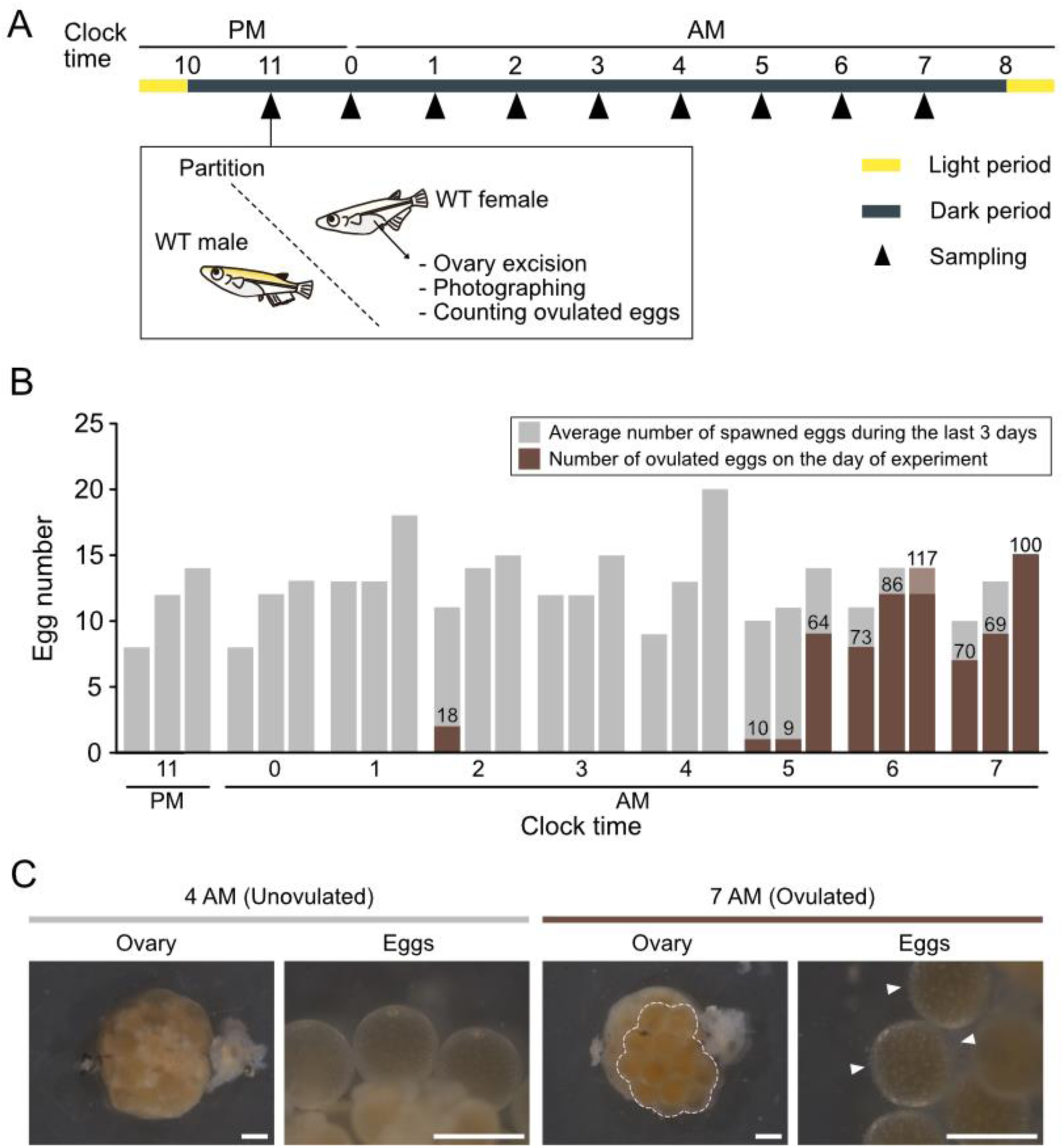
Ovulation occurs around 2 hours before the lights-on. (A) Schematic diagram showing the experimental design. (B) Bar graph indicating the average number of spawned eggs during the last three days before the experiment (gray) and number of ovulated eggs on the day of experiment (brown), with the clock time on the horizontal axis and the egg number on the vertical axis. Each time bin on the horizontal axis includes three bars, each of which represents the data for one individual. The numbers in the graph indicate the percentage (%) of the number of ovulated eggs on the day of experiment relative to the average number of spawned eggs during the last three days. (C) Representative photographs of the gross anatomy of the ovary and eggs viewed from the dorsal side. Arrowheads indicate the fibrous membranes, which characterize the ovulated eggs. Scale bars represent 1 mm.

### Intact medaka pairs perform sexual behavior after ovulation even in the dark

We further scrutinized the timeline of sexual behavior under the natural conditions by video-recording of intact WT pairs between 4:30 AM – 9:00 AM. In this experiment, the male and female pairs were not separated with a partition and were allowed to interact freely with each other, which is somewhat different from the conditions for sexual behavior analyses of KO females (Figure 1). We used LED lights that emit 940 nm wavelength infrared light for the behavioral recording during the dark period (4:30-8:00 AM). We confirmed that this infrared LED lights are invisible to medaka (Figure S2, methods are described in Document S2). We found that the pairs started to perform spawning approximately after 6:30 AM (Figure 3A), when it is suggested that ovulation has already started (see Figure 2). This result suggests that the motivation for sexual behavior is already increased after the ovulation, which is consistent with the results of *gnrh*^-/-^ and *lhb*^-/-^ females; the occurrence of sexual behavior is dependent on the presence of ovulation (Figure 1).

**Figure 3.**
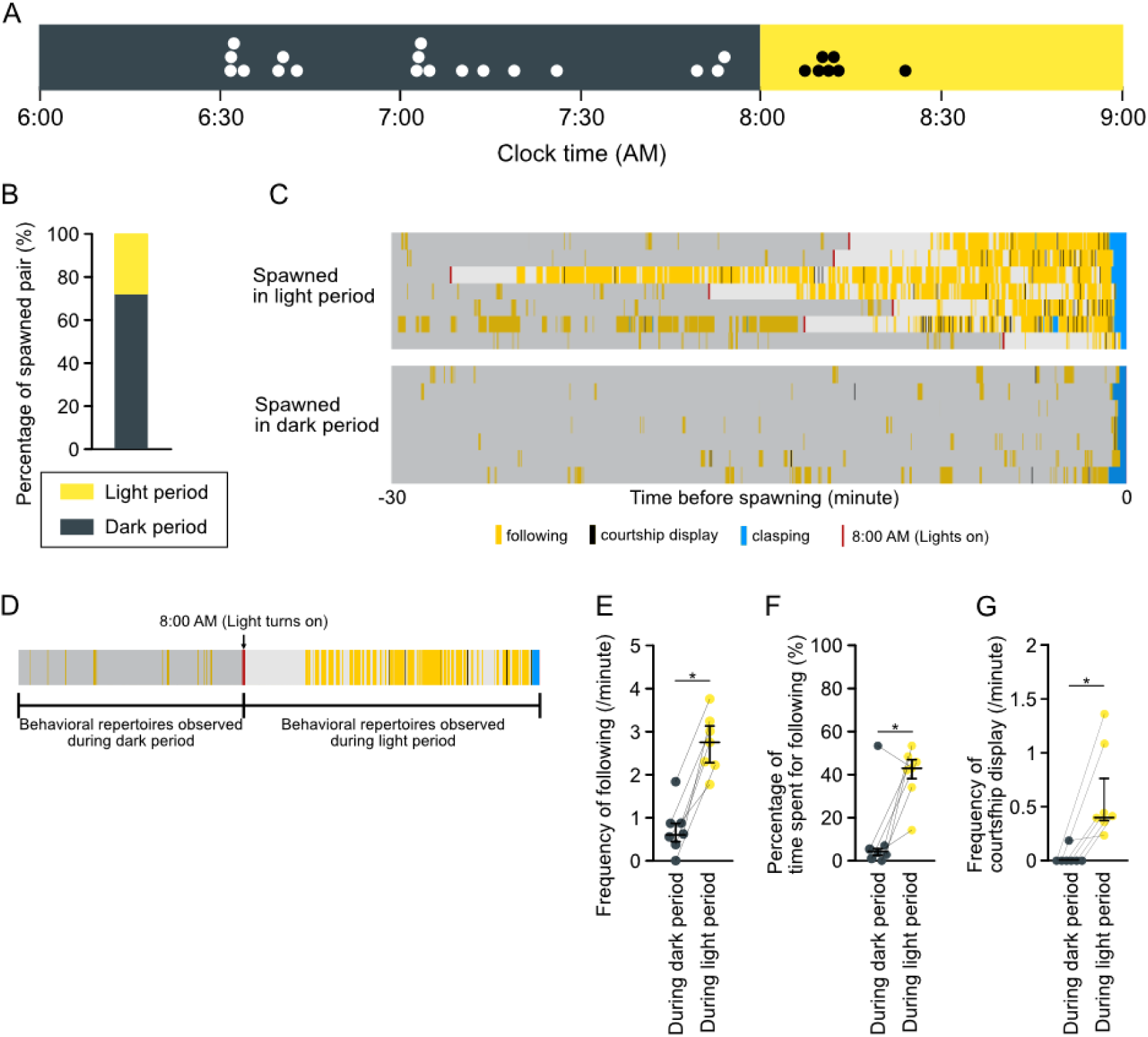
Intact pair can spawn before the light-on. Sexual behavior of the WT pair was analyzed between 6:00 AM - 9:00 AM. Infrared LED lights (940 nm wavelength) stayed on until 8:00 AM, and then the visible lights were turned on. (A) The time course of the occurrence of spawning in the dark (black bar, 6:00 AM - 8:00 AM) and in the light (yellow, 8:00 AM - 9:00 AM). Each dot indicates a different pair. (B) The percentage of spawned pair during the dark period (black) and the light period (yellow). (C) The time course (the time of spawning set to zero) of sexual behavior repertoires during 30 minutes before spawning is shown as raster plots. Raster plots of the pairs that spawned during the dark period shown in this figure (7 pairs) were randomly chosen from 18 pairs. (D) Representative raster plot showing the difference between dark and light period s in the occurrence of behavioral repertoires before clasping in (E-G). (E-G) Scatter and whisker plots indicating frequency of following trial (E) (\**P* = 0.01563), percentage of the time spent for following (F) (\**P* = 0.03125), and frequency of courtship display (G) (\**P* = 0.01563).

Interestingly, 7 of 25 pairs (28%) spawned during the light period (8:00-9:00 AM), whereas 18 of 25 pairs (72%) spawned during the dark period (4:30-8:00 AM) (Figure 3B). This result indicates that medaka pairs perform sexual behavior even during the dark period. We therefore compared the sexual behavior repertoires performed during the light and dark periods (Figure 3C). We found that the males during the light period frequently showed following and courtship display prior to clasping and subsequent spawning (Video S1). In contrast, they rarely showed following and courtship display during the dark period, although they finally show clasping and spawning (Figure 3C and Video S2). We also compared male following and courtship display during dark and light periods performed by each pair that culminated in spawning in the light period (Figure 3D). Although following and courtship display were observed even in the dark period, the frequency of following (Figure 3E), the percentage of time spent for following (Figure 3F), and the frequency of courtship display (Figure 3G) during the dark period were significantly lower than those during the light period. These results suggest that the male sexual behavior repertoires dependent on the visual cues (following and courtship display) cannot be easily performed during the dark period, while the other repertoires not dependent on the visual cues (clasping and spawning) are rather normally performed. Since males switch patterns of sexual behavior repertoires depending on the light conditions, females may receive different courtship cues from males under the dark and light conditions.

### Progesterone administration does not induce ovulation but partially reinstates the sexual receptivity in *lhb*^-/-^ females

The behavioral analyses thus far show that the occurrence of ovulation facilitates the female sexual receptivity, suggesting that certain substances associated with ovulation may affect female receptivity. Since the ovulatory cycle is accompanied by fluctuations of some kinds of sex steroid hormones^27^, we hypothesized that some of these sex steroid hormones may play important roles in the activation of sexual receptivity of female medaka. Among them, we focused on the progesterone derivatives, which is known to be released in strong synchrony with ovulatory cycles and has been suggested to be involved in female receptivity in mammals^28,29^. We administered progesterone to *lhb*^-/-^ females by water exposure and analyzed their sexual behavior with WT male (Figure 4A). One day after starting progesterone administration, three out of six pairs of *lhb*^-/-^ females with progesterone administration exhibited clasping. Furthermore, another pair of *lhb*^-/-^ females performed clasping two days after starting progesterone administration (Figure 4B). It should be noted that clasping was observed 25-75 minutes after the female and male started interactions (Figure 4C), showing that the latency to the first clasping is prolonged compared with the natural clasping performed by WT pair (2-30 minutes, see Figure 1). Although clasping was reinstated by progesterone administration, spawning was not accompanied with clasping in this experiment. To examine whether progesterone-administered female possesses normal ovary or not, we analyzed the morphology of ovaries from both 10-day progesterone-administered and control *lhb*^-/-^ females. We did not find any ovulated eggs in the 10-day progesterone-administered ovaries of *lhb*^-/-^ females or in the DMSO-administered control females (Figure 4D). Similarly, there was no discernible difference in the ovarian histology between the 10-day progesterone-administered and the control *lhb*^-/-^ females (Figure 4E). These results suggest that the administered progesterone did not induce ovulation but may have affected certain neurons in the brain to activate female sexual receptivity.

**Figure 4.**
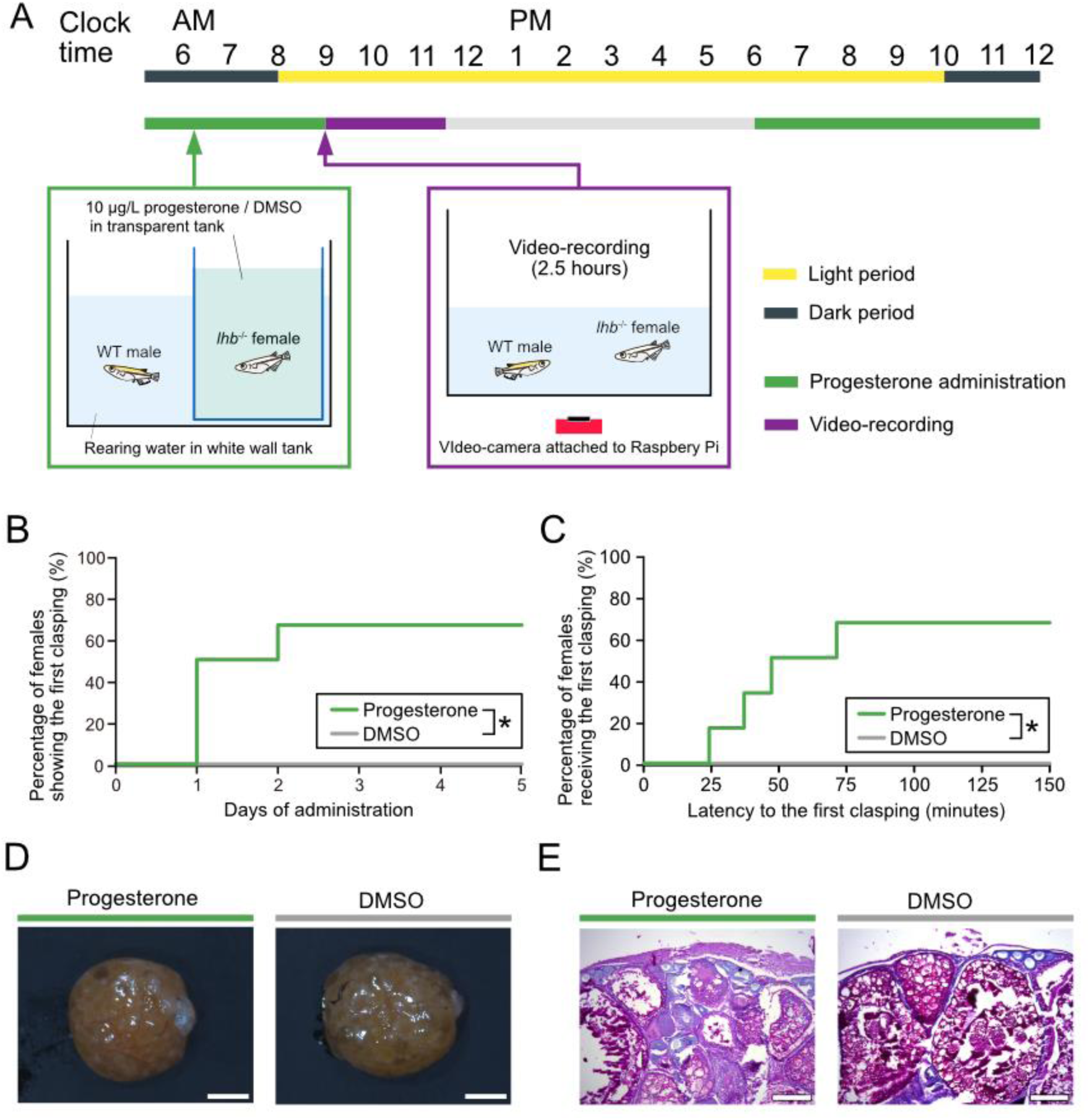
Progesterone administration reinstates female receptivity in *lhb*^-/-^ females without inducing <u>ovulation</u>. (A) Schematic diagram showing the procedure of progesterone administration. Progesterone was administered from 6:00 PM to 9:00 AM by water exposure. At the start of video recording (9:00 AM), *lhb*^-/-^ female was moved to a recording tank and paired with a intact WT male. Video recording was performed for 2.5 hours. (B) Reverse Kaplan-Meier curves indicating the percentage of females showing clasping after progesterone administration (n = 6 pairs for each treatment, \**P* = 0.02 by log-rank test). (C) Reverse Kaplan-Meier plots indicating the percentage of females receiving the first clasping (n = 6 pairs for each treatment, \**P* = 0.02 by log-rank test). The data were obtained from the video recorded three days after the progesterone administration. (D) Representative photographs of the dorsal views of the ovaries of progesterone administered/control females. Scale bars: 3 mm. (E) Representative photomicrographs of hematoxylin/eosin-stained sections of the ovaries of progesterone administered/control females. Scale bars: 200 µm.

### Progesterone receptors are expressed in various brain regions of the female

To examine the actual distributions of neurons expressing progesterone receptors, which may potentially mediate sexual receptivity of the female medaka, we examined the localization of neurons expressing *progesterone receptor* (*pgr*), which has been suggested to be involved in the sexual behavior in some rodent species^30,31,32^. *In situ* hybridization analysis revealed that *pgr* was expressed in the area ventralis telencephali pars supracommissuralis (Vs), the area ventralis telencephali pars posterior (Vp), the area preoptica (POA), the nucleus preopticus pars parvocellularis (POp), the nucleus preopticus pars magnocellularis (POm), the nucleus posterioris periventricularis (NPPv), and the nucleus ventral tuberis (NVT) of the female brain (Figure 5), which are considered to be involved in the regulation of sexual behavior in teleosts^33,34,35,36^.

**Figure 5.**
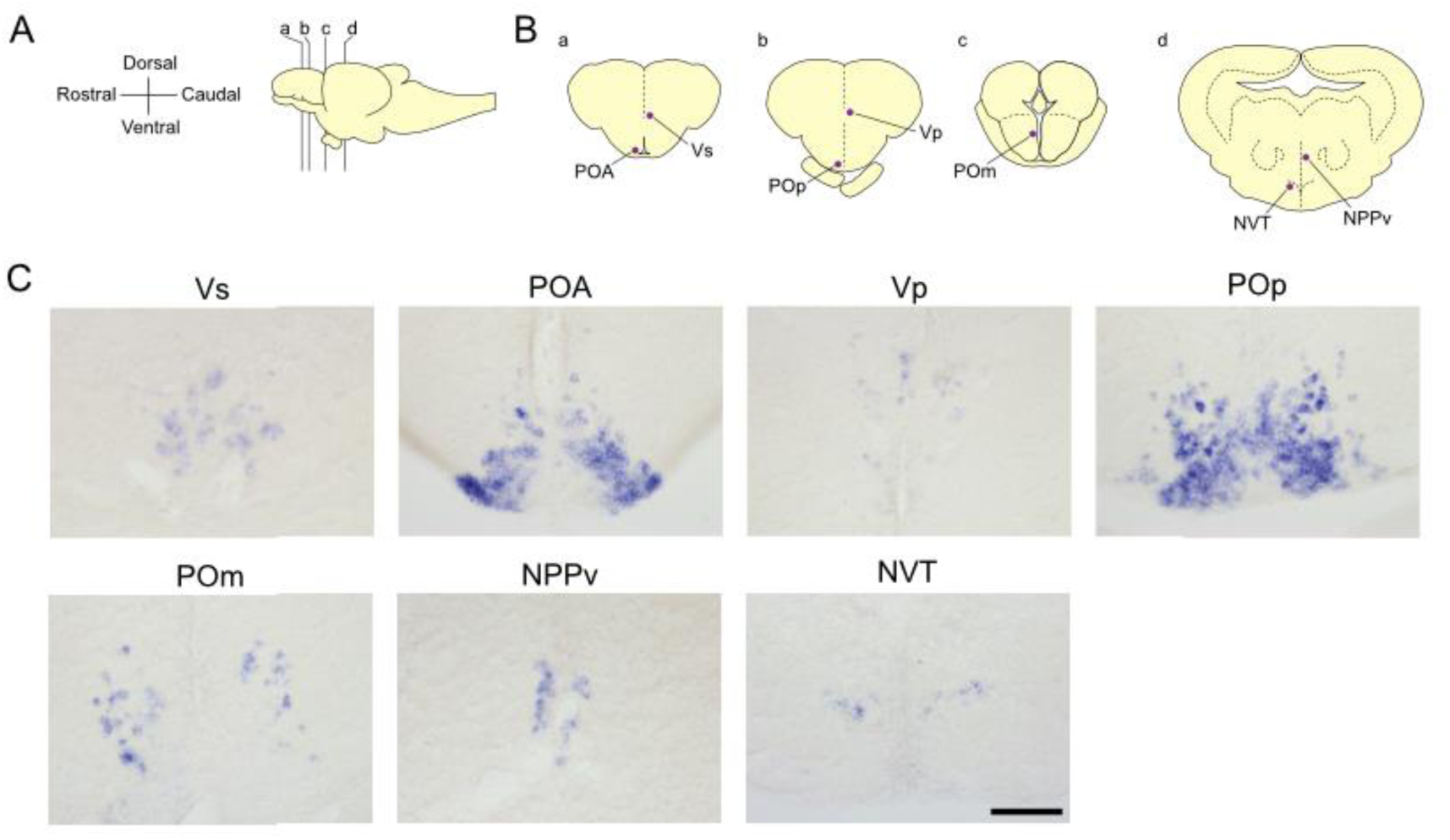
*pgr* is expressed in some brain nuclei of female medaka. (A) Illustration of the lateral view (rostral to the left) of medaka brain showing the levels of the frontal sections in (B). (B) Illustrations of the frontal sections showing the localization of neurons expressing *pgr* (purple signals); Vs, POA, Vp, POp, POm, NPPv, and NVT. (C) Representative photomicrographs of ISH for *pgr* in some brain nuclei. Scale bars: 50 µm.

## Discussion

### Ovulation plays an important role in the facilitation of female sexual receptivity

In the present study, we analyzed the sexual behavior of the anovulatory female medaka in which one of the HPG-axis related genes was knocked-out (*fshb*^-/-^/*lhb*^-/-^/*gnrh*^-/-^). We found that the anovulatory female medaka failed to exhibit the male-receptive behavior (Figure 1). Furthermore, analyses of the temporal relationship between ovulation and activation of female sexual receptivity revealed that clasping is performed just after the female ovulation (after 6:30 AM (1.5 hours before the lights-on), Figure 3A). Considering the result that ovulation occurs around 5-7 AM (1-3 hours before the lights-on, Figure 2B), which is consistent with the previous studies^37,38^, sexual behavior is performed approximately 1-2 hours after the female ovulation. These results suggest that ovulation plays critical roles in the activation of female receptivity. Furthermore, administration of progesterone, which is known to be released in strong synchrony with ovulatory cycles, partially reinstated the female receptivity in *lhb*^-/-^ female without inducing ovulation (Figure 4). Histological analysis demonstrated that progesterone-receptor is expressed in some brain nuclei that have been considered to be involved in the control of sexual behavior ^33,34,35,36^ (Figure 5). Taken together, we suggest that certain substance(s) released from the ovaries during ovulation, such as progesterone, plays and important role in the activation of female sexual behavior, especially the female receptivity to be clasped by male.

In most of teleost species, sexual behavior is usually performed by the individuals having mature gonads and gametes. For example, seasonal breeders show sexual maturation and sexual behaviors during the breeding season, which is a phenomenon that reflects such synchronization of gonadal state and sexual behavior. It should be noted that females not only perform gametogenesis but also show ovulation during this season for successful reproduction. However, the mechanism of synchronization of gonadal state and behavior has remained elusive. Medaka has been recognized as a seasonal breeder, and its breeding condition can be experimentally controlled by photoperiod^37^. Under the long-day photoperiod such as that used in the present study (14L and 10D, see Methods), the male and female medaka are in the breeding condition, and they possess mature gonads and spawn daily. On the other hand, they become non-breeding under the short-day photoperiod (10L and 14D), even when the water temperature is maintained at breeding condition^39^. Here, we analyzed the sexual behavior of *fshb*^-/-^ female medaka under the breeding condition and found that they do not exhibit sexually receptive behavior despite the normal courtship display by male (Figure 1). In addition, neither *lhb*^-/-^ nor *gnrh*^-/-^ females, which do not ovulate, showed clasping. These results clearly show the importance of gonadal maturation, especially ovulation, for the activation of female sexual receptivity. Interestingly, males normally courted to *fshb*^-/-^/ *lhb*^-/-^/*gnrh*^-/-^ female, which indicates that successful reproduction is highly dependent on the female sexual receptivity. These behaviors are considered to be one of the examples of “female choice” ^40^.

### Involvement of sex steroid hormones mediating ovulatory cycles and female receptivity in teleosts

There are not a few studies in mammals that analyzed the relationship between the ovulatory cycle and the female sexual behavior. In mice and rats, female receptivity has been known to be mediated by sex steroid hormones, the levels of which fluctuate in synchrony with the ovulatory cycle. In fact, it has been suggested that one of the sex steroid hormones, estrogen, plays a key role in synchronizing female receptivity with ovulation; estrogen implantation upregulates the lordosis behavior^41^, whereas gene KO of estrogen-receptor (Esr) disrupts it^42^, suggesting the involvement of estrogen/Esr signaling in the female sexual receptivity. Recent studies have shown that Esr-expressing neurons in the ventromedial hypothalamus (VMH) change their gene expression and synaptic connectivities, which promote the female sexual receptivity^43,44,45^. Also, it has been demonstrated that Esr-expressing neurons that play key roles in ovulation also project to the brain regions controlling the sexual behavior. In female mice, for example, kisspeptin neurons in the anteroventral periventricular nucleus (AVPV^Kiss^ neurons), which is known to be sensitive to estrogen and triggers GnRH surge by projecting to the GnRH neurons, have also been reported to project to the ventrolateral subpopulation of VMH neurons and promote lordosis behavior^46^. Thus, estrogen serves as a mediator of ovulatory cycles and female receptivity. In addition to estrogen, progesterone has also been considered as a sex steroid hormone responsible for the upregulation of female receptivity. Progesterone implantation into the VMH facilitates female receptivity accompanied by lordosis in rats^47,48^. Recent studies have reported that specific population of *pgr*-expressing neurons in the VMH are essential for the female sexual behavior in mice^31,49^. Thus, female sexual receptivity is regulated by the neurons that directly sense periodic fluctuations of sex steroid hormones synchronized with the ovulatory cycles. On the other hand, the kisspeptin system in non-mammalian species has been shown not to be involved in the HPG axis regulation^8,12^, and recent studies have unraveled that there are other mechanisms for the HPG axis regulation in non-mammalian species^50,51^. However, it has not been studied whether such coordinated regulations of the HPG axis and the sexual behaviors via sex steroids also exist in the non-mammalian species as well.

Recent studies in medaka have demonstrated various kinds of functions of sex steroid hormones in the brain. Estrogen has various neuromodulatory functions including the HPG-axis regulation ^52,53,54,55,56,57^. In addition, some *Esr*-expressing neurons play pivotal roles in the activation of female receptivity. Neurons in the ventral telencephalon and/or preoptic area that express *esr2b*, one of the subtypes of Esr genes in teleosts, mediate female receptivity probably by releasing neuropeptides^58,59^. However, since *esr2b* KO female medaka normally ovulates, it remains unclear whether ovulatory cycle is involved in the control of female sexual behavior. In the present study, we demonstrated that ovulation is essential for facilitating female sexual receptivity, and progesterone administration partially reinstates the female receptive behavior in *lhb*^-/-^ female without inducing ovulation. The present result that progesterone administration did not induce ovulation is consistent with the previous study showing that progesterone administration induced oocyte maturation but not ovulation in the hypophysectomized females^25^. Thus, our present results suggest that female receptivity is upregulated in synchrony with ovulatory cycles via some progesterone signaling. Considering the results that clasping is performed 1-2 hour after the ovulation, we hypothesized that progesterone exerts slow/chronic effects rather than acute ones and focused on the nuclear progesterone receptor Pgr instead of the membrane progesterone receptor. Histological analyses revealed that *pgr* is expressed in some brain regions that are supposed to be involved in the activation of sexual behavior in teleost species^33,34,35,36^. Thus, our present behavioral and histological results suggest that *pgr*-expressing neurons may somehow modulate the neural circuit properties of relevant brain regions around the time of ovulation, which may mediate female sexual receptivity. Although the action of progesterone is conceivably received by *pgr*-expressing neurons and contributes to the facilitation of female receptivity, the mechanisms of progesterone actions in medaka have not been clearly understood. Some studies using other teleosts reported that blood progesterone level fluctuates around the timing of ovulation and shows the highest level just before the spawning^60,61^. Future studies should analyze the relationship between the dynamics of progesterone levels and the female sexual receptivity and its neural mechanisms.

Another possible candidate for the ligands that bind to Pgr expressed in the brain may be 17α,20β-dihydroxy-4-pregnen-3-one (17α,20β-DHP). Accumulating evidence strongly suggests that 17α,20β-DHP is released in a surge mode from the ovary, which mainly contributes to induce the final oocyte maturation and subsequently ovulation in some teleost species including medaka^62,63,64,65,66^. It has also been reported that *pgr* serves as a receptor for 17α,20β-DHP^27,67^. Considering the fact that 17α,20β-DHP is released about 14 hours after the lights-on (around 22:00 PM)^25,66,68^, it is possible that 17α,20β-DHP may activate *pgr*-expressing neurons in a chronic manner to modulate neural circuits controlling female sexual receptivity. It should be noted that not of all the females in the present study showed clasping by progesterone administration (Figure 4B), and the latency to the first clasping was rather long even in the pairs that exhibited clasping (Figure 4C). One possible explanation is that the present pharmacological treatment may not have fully mimicked the progesterone signaling in the intact female. Although *in vitro* analysis in medaka revealed that progesterone binding to Pgr leads to an increased level of transcriptional activity, it is weaker than that of 17α,20β-DHP binding to Pgr^69^. Also, the progesterone concentration is extremely low compared with 17α,20β-DHP, and it also remains unclear whether progesterone concentration fluctuates diurnally synchronized with the ovulatory cycle. In contrast, 17α,20β-DHP is a ligand for Pgr, which is known to fluctuate diurnally. Unfortunately, however, the administration of 17α,20β-DHP can induce ovulation and make it difficult to test the possible function of Pgr signaling in the brain. Therefore, in the present study, progesterone was administered during the time window from 6:00 PM to 9:00 AM, mirroring the diurnal fluctuations observed in 17α,20β-DHP. Another plausible explanation may be that factors other than 17α,20β-DHP/Pgr signaling are responsible for the female receptivity but were lost simultaneously with the deficiency of ovulation in *lhb*^-/-^ female. Interestingly, Fleming *et al.*, 2022 showed that prostaglandin E2 (PGE2) is involved in the activation of female receptivity via Ptger4b-expressing neurons^70^. Since PGE2 is thought to be acutely released around the timing of ovulation^70^, it may be possible that multiple factors cooperatively modulate the activity of relevant neurons and facilitate fine-tuning of female receptivity.

### Medaka can spawn before ‘dawn’ without visual cues as long as the female ovulates

It has been generally considered that visual cue is one of the most important sensory cues for triggering various behaviors of medaka including sexual behaviors^71,72,73^, and medaka have been supposed to perform sexual behavior soon after the lights-on depending on the visual cues. On the other hand, it has also been reported in medaka under the natural condition that they spawn before sunrise^74^. Thus, it has been rather controversial whether medaka perform sexual behavior even during the dark period. In the present study, we observed the sexual behavior of WT medaka pair by using an infra-red wavelength video-recording system and discovered that WT medaka pairs do perform sexual behavior approximately from 6:30 AM. Since the lights were set to turn on at 8:00 AM in the present experiment, it follows that medaka pairs can perform sexual behavior even during the dark period, which is equivalent to the dawn under the natural conditions. We also found out that frequency of the courtship display, which is a stereotypical courtship behavior performed by male, was significantly decreased in the dark period in comparison with the light period (Figure 3G). It suggests that the male courtship display is highly dependent on the visual cues. It also suggest that the male medaka perform clasping and spawning partly dependent on the other sensory cues such as the olfactory and/or somatosensory cues, which is consistent with the previous study; occlusion of the nasal cavities of male medaka significantly decreased the frequency of courtship behavior and disrupted clasping^75^. Since there are some studies using other teleost species showing that the male courtship behaviors are triggered by female pheromonal substances^76,77,78^, it is also possible that similar pheromonal stimuli may also trigger male courtship behavior in medaka.

Since the female medaka allowed males to wrap their body and perform spawning without receiving the courtship display from the male, it follows that the female medaka do not need to receive the male courtship display as a visual information at least in the dark. From the fact that sexually receptive behavior of the female was observed after the start of ovulation without receiving the courtship display from males, it is suggested that the ovulatory signals, perhaps mediated by *pgr*-expressing neurons, is involved in the facilitation of female sexual receptivity but not in the modulation of sensory systems for receiving various cues from male.

In summary, our present study provides insights into the neuroendocrine mechanism that synchronize the timing of sexual behavior with the ovulatory cycles in teleost species, which contributes to the successful reproduction.

## Acknowledgements

We thank Mses. Hiroko Tsukamoto and Hisako Kohno (The University of Tokyo) and Erina Fukunaga (Nagahama-Institute of Bio-Science and Technology) for excellent care of the fish used in this study. We are also grateful to Dr. Kana Ikegami for helpful discussion and comments.

Financial support: This work was supported by JSPS KAKENHI Grant Numbers JP19J21828 (to S.T.), JP2201130 (to S.T.), JP22KJ2986 (to S.T.), JP21K15140 (to M.N.), JP26221104 (to Y.O.), JP21K06262 (to Y.O.), and JP20H03071 (to C.U.). This work was also supported by World-leading Innovative Graduate Study Program for Life Science and Technology (WINGS-LST), the University of Tokyo WISE Program (Doctoral Program for World-leading Innovative & Smart Education), MEXT, Japan (to S.T.).

## Author contributions

Conceptualization: S.T., M.N. and C.U.; Validation, Formal Analysis, Data Curation, and Visualization: R.S. and S.T.; Investigation: S.T., R.S., M.N., and C.U.; Project Administration: S.T., M.N., and C.U.; Resources, and Supervision: Y.O. and C.U.; Writing – Original Draft: S.T.; Writing – Review & Editing: S.T., R.S., M.N., C.U., and Y.O.; Funding Acquisition: S.T., M.N., C.U., and Y.O.

## Materials & Methods

### Animals

Male and female d-rR wild type (WT) medaka (*Oryzias latipes*) and gene-knockout (KO) lines of *fshb* (*fshb*^+/+^, *fshb*^+/-^ and *fshb*^-/-^), *gnrh1* (*gnrh1*^+/+^, *gnrh1*^+/-^ and *gnrh1*^-/-^). and *lhb* (*lhb*^+/+^, *lhb*^+/-^ and *lhb*^-/-^), which were generated in Takahashi *et al.*, 2016^17^, were used in the present study. All fish were maintained in shoals with water circulation (Labreed, IWAKI Co., Ltd., Tokyo, Japan) under 14-hour light/10-hour dark photoperiod (lights-on at 8:00 AM and -off at 10:00 PM) condition at a water temperature of 27 ± 2 °C. They were fed with live brine shrimp and/or commercial flake food (Otohime; San-u fish farm, Osaka, Japan) for three or four times per day. All experiments were performed using sexually mature 3-7 months old males and females. The fish maintenance and the experiments were conducted in accordance with the protocols approved by the Animal Care and Use Committee of the University of Tokyo (permission number 20-6) and the Animal Experiment Committee of the Nagahama Institute of Bio-Science and Technology (approval number: 098).

### Method details

#### Genotyping and screening of KO medaka

Since homozygous KO females of *fshb*, *gnrh1*, and *lhb* (*fshb*^-/-^, *gnrh1*^-/-^ and *lhb*^-/-^) are infertile^17^, the KO medaka lines were obtained from heterozygous males and females, and their siblings were used for phenotypic studies. We extracted genomic DNA from clipped tail fin and used it as a template for real-time PCR, and amplified sequence containing each mutation site by using primers described in the previous study^17^. The temperature profile of the reaction was 95 °C for 150 seconds of pre-denaturation, and 40 cycles of denaturation at 95 °C for 10 seconds, annealing at 60 °C for 10 seconds, and extension at 72 °C for 15 seconds. We verified the genotype by using melting curve of the PCR products generated by the following temperature profile: 95 °C for 10 seconds, 65 °C for 10 seconds, and 97 °C for 1 second.

#### Recording setup for sexual behavior analysis

Analyses of sexual behavior of medaka were performed under the same photoperiod and temperature conditions for fish maintenance. Behavioral recordings were conducted under two conditions as described below.

Condition A: As described in our previous study^19^, male and female pairs were placed in tanks with white walls and transparent square bottom (15.0 × 15.0 cm), and were separately kept until recording by putting them in a tank with transparent perforated partition diagonally across the tank. The behavioral recording was performed by using Raspberry Pi ZERO W (Raspberry Pi Foundation, Cambridge, UK) equipped with Raspberry Pi Module Camera Module V2.1 (Raspberry Pi Foundation) device installed with a software for automated recording developed in our previous study^19^. For making the same recording background, we placed the thin white paper in each tank.

Condition B: Male and female pairs were placed in tanks with black bottom (13.5 × 22.5 cm), three black walls and one transparent wall for recording. The water depth was maintained at ∼ 12 cm. Behavioral recording was performed by using Raspberry Pi 3 Model B (Raspberry Pi Foundation) device equipped with Raspberry PiNoir Camera Module V2.1 (Raspberry Pi Foundation) and installed with the same software with Condition A.

#### Sexual behavior analysis of KO females

For analyzing the sexual behavior of KO female, we performed behavioral analysis of condition A, the same method as described in our previous study^19^. We paired KO female with WT sibling male and placed it in the experimental white tank three days before behavioral recording. At 6-7 PM of the day before recording, females and males were separated by putting a transparent perforated partition diagonally across the tank. At 9:00 AM (one hour after the onset of the light period) of the following day, the partition was removed, and video recording was conducted for one hour by running a software for automated recording developed in our previous study^19^.

After obtaining behavioral videos, we identified the behavioral repertoires and annotated four behavioral repertoires during sexual behavior; “following”, “courtship display”, “clasping”, and “spawning” by using *Ethogramer*^19^, while replaying the first 30 minutes of the videos from the time the partition was removed, and male and female started to interact. Each repertoire was referred to as “approaching to courting orientation”, “head-up I to courting round dance”, “head up II to copulation”, and “spawning”, respectively in the previous literature^20,21^ (See also Video. S1). After behavioral annotation, we calculated the following behavioral parameters as described below:

- Frequency of following (/minute)
- Percentage of time spent for following (%)
- Frequency of courtship display (/minute)
- Latency to the first clasping (minute)
- Latency to spawning (minute)

It should be noted that the frequency data was calculated for the entire analysis period (30 minutes), and the latency was regarded as 30 minutes when the pairs did not show clasping and/or spawning during the analysis period.

#### Ovarian morphology

We used WT females that were confirmed to spawn more than seven fertilized eggs for three consecutive days after pairing with WT male (female gonadosomatic index (gonadal weight/body weight × 100 (%)): 11.8 ∼ 20.8). During 11:00 PM – 7:00 AM, three ovaries were dissected out every hour, observed and photographed under a Leica S9D (Leica Microsystems, Wetzlar, Germany) stereomicroscope equipped with a Leica MC120HD (Leica Microsystems) digital camera.

We observed the gross morphology of the ovaries, then picked up the oocytes and eggs by tearing open the ovarian surface epithelium. We distinguished ovulated eggs judging from their size, color, and fibrous membrane structure. The timing of ovulation was estimated by comparing the number of ovulated eggs and the average number of spawned eggs for last three days, which was defined as the number of eggs that were attached to the female abdomen.

#### Sexual behavior analysis of WT pairs before and after the lights-on

WT female and male pairs that were confirmed to spawn consecutively were placed in the condition B experimental tank. Video recording was performed during 4:30 - 9:00 AM (4:30 - 8:00: dark, 8:00-9:00 light) under the 940 nm wavelength light by using an LED unit (FRS5JS; Optosupply, Hong Kong, China). It should be noted that 940 nm wavelength light was confirmed by optomotor response (OMR) experiment (Figure S1) to be invisible for medaka.

We detected the timing of spawning and annotated the behavioral repertoires that were performed during 30 minutes before spawning by using *Ethogramer*. We compared the following three parameters of the pairs that spawned during the period from dark to light (Figure 3D).

- Frequency of following (/minute)
- Percentage of time spent for following (%)
- Frequency of courtship display (/minute)

Fertilization rate was calculated by the same method as described above.

#### Progesterone administration and gonadal analyses

We prepared a transparent tank (7.4 × 14.0 × 11.2 cm), which can be put into the observation tank used in recording condition A, for progesterone or DMSO administration only to the female, while the female was visible to the male in the observation tank. At 6:00 PM, 10 µL of 1 g/L progesterone (Wako, Osaka, Japan) in Dimethyl sulfoxide (DMSO, FUJIFILM Wako Pure Chemical, Osaka, Japan) was dissolved in 1 L rearing water (final concentration of progesterone: 10 µg/L). For the control, the same volume (10 µL) of DMSO was dissolved in the rearing water. Either progesterone or DMSO water was poured in the transparent tank, and 1 L of rearing water was also poured in the observation tank (Fig.4A). *lhb*^-/-^ female and WT male were placed in the transparent and white tanks, respectively. At 9:00 AM, *lhb*^-/-^ female was transferred from the transparent tank to the behavioral observation tank, and the transparent tank was removed. Consequently, the *lhb*^-/-^ females were exposed to progesterone only during 6:00 PM to 9:00 AM (15 hours) (Fig.4A). Behavioral recording was conducted between 9:00 - 11:30 AM, and this experiment was repeated for five days. We counted the days of progesterone administration until they showed the first clasping, and further analyzed the latency to the first clasping on the third day after the start of progesterone administration.

Ten days after the progesterone exposure, *lhb*^-/-^ females were deeply anesthetized with 0.02% MS222, and their ovaries were dissected out. The gross morphology of the ovaries was photographed from the dorsal side. Then the ovaries were immediately fixed overnight with Bouin’s fixative (picric acid saturated solution : formaldehyde : acetic acid = 15 : 5 : 1, all chemicals were sourced from FUJIFILM Wako Pure Chemical Corporation, Tokyo, Japan) at 4 °C. The fixed ovaries were dehydrated with ethanol, cleared with xylene, and embedded in paraffin (Histprep568, FUJIFILM Wako Pure Chemical Corporation). The embedded specimens were sectioned serially at a thickness of 5 μm and were stained with hematoxylin and eosin (HE) (FUJIFILM Wako Pure Chemical Corporation. Photo images were acquired using DM5000B (Leica Microsystems) equipped with DFC310 FX (Leica Microsystems).

#### *In situ* hybridization

*In situ* hybridization was performed as described in the previous study ^22^. WT females were deeply anesthetized with 0.02% MS222 and quickly fixed by perfusion with 4% paraformaldehyde (PFA) (Nakarai Tesque, Kyoto, Japan) in 1.0 × PBS (Takara Bio) for 30 seconds from the bulbus arteriosus using a borosilicate glass pipette made from capillaries with 1.5-mm outer diameters (GD-1.5; Narishige, Tokyo, Japan). After perfusion, we decapitated the fish, dissected out the whole brain and soaked it in a fixative containing 4% PFA in 1.0 × PBS for 10 minutes at room temperature. We substituted it with 30 % (w/v) sucrose (FUJIFILM Wako Pure Chemical Corporation) in 1.0 × PBS overnight at 4 °C. The brain was embedded in 5% ultra-low gelling agarose (Sigma-Aldrich, Darmstadt, Germany) / 20% sucrose (w/v) in 1.0 × PBS, and serial frontal cryosections were cut at 25 μm intervals by using a cryostat (CM 3050S; Leica Microsystems). The sections were mounted onto CREST-coated glass slides (Matsunami, Osaka, Japan). The sections were treated with 1 μg/mL proteinase K in 1.0 × PBS for 15 minutes at 37 °C, postfixed with 4% PFA in 1.0 × PBS for 10 minutes, treated with 2 mg/mL glycine (FUJIFILM Wako Pure Chemical Corporation) in 1.0 × PBS for 5 minutes, and then incubated with 0.25% acetic anhydride (FUJIFILM Wako Pure Chemical Corporation) in 0.1 M triethanolamine (Sigma-Aldrich) for 10 minutes. The sections were subsequently hybridized overnight with digoxigenin (DIG)-labeled RNA probe for *pgr*, which was prepared according to Zempo *et al.*, 2018 ^23^, at 58 °C. Hybridization signals were detected using alkaline phosphatase-conjugated anti-DIG antibody (Roche Diagnostics, Basel, Switzerland) and 5- nitro blue tetrazolium/bromo-4-chloro-3-indolyl phosphate (NBT/BCIP) (Roche) as chromogenic substrates. Photo images were acquired using the same equipment used in HE-staining of ovary. We used the nomenclature of the medaka brain nuclei, as described in the previous study^24^.

## Statistical analysis

Statistical analysis and graph drawing were performed by using R (R Core Team 2020). Mann– Whitney *U* test was performed for comparison between WT and KO, which were shown by the whisker and scatter plot, except for the latency to the first following/spawning. For the latency to the first following/spawning and spawning days after progesterone exposure in Figure 4, Kaplan–Meier plots were drawn, and the differences between each Kaplan–Meier curve were tested for statistical significance using Log-Rank test. For comparison of the behavioral parameters between dark period and light period (Figure 3), Wilcoxson signed-rank sum tests were conducted and shown by the whisker and scatter plot. A *P*-value less than 0.05 was considered statistically significant.

