## Supplementary Information for "Sexual receptivity increases in synchrony with the ovulatory cycle in female medaka"

**Figure S1**

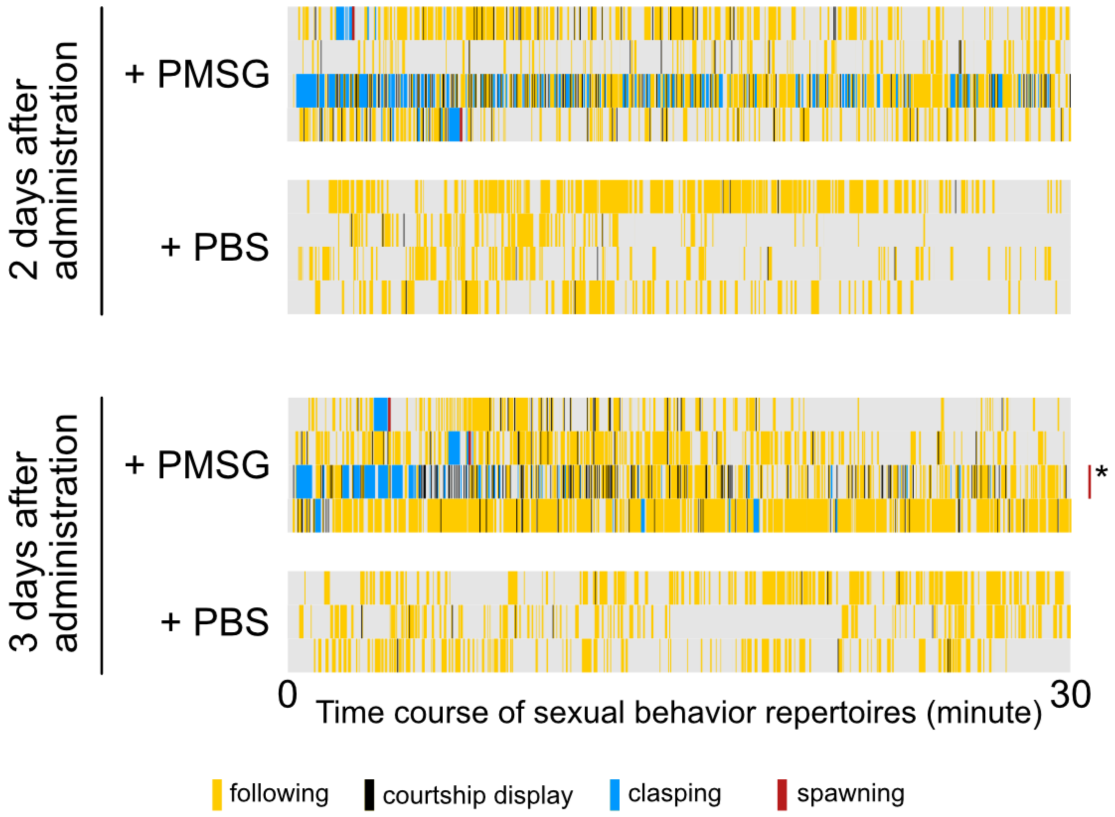
2 days after administration
+ PMSG
+ PBS
3 days after administration
+ PMSG
*
+ PBS
0
Time course of sexual behavior repertoires (minute)
30
following
courtship display
clasping
spawning

Supplementary Figure S1. PMSG administration reinstates clasping in *gnrh1* KO female (Related to Figure 1)

Time course (during 30 min) of the sexual behavior repertoires 2 or 3 days after administration shown as raster plots. +PMSG raster plots show the behavioral transition of the pairs in which females were administrated PMSG, and +PBS raster plots show those in which females were administrated PBS as a negative control. The red bar with an asterisk indicates that this pair showed spawning 30 minutes after the behavioral analysis.

**Figure 2**

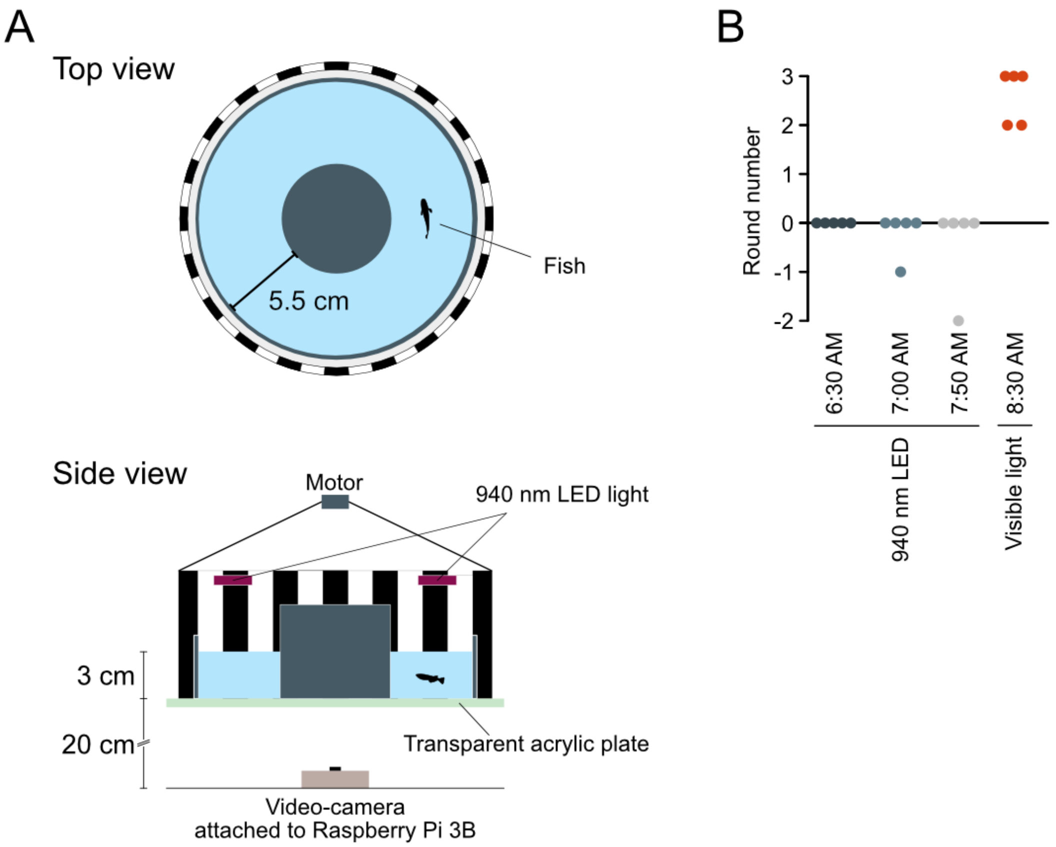
A
Top view
5.5 cm
Fish
Side view
Motor
940 nm LED light
3 cm
20 cm
Transparent acrylic plate
Video-camera
attached to Raspberry Pi 3B
B
Round number
3
2
1
0
-1
-2
6:30 AM
7:00 AM
7:50 AM
8:30 AM
940 nm LED
Visible light

Supplementary Figure S2. Infrared wavelength light is invisible to medaka (Related to Figure 3)

(A) Schematic diagram of the setup for optomotor response experiment. Black-and-white paper cylinder is rotated by a motor control unit attached to Rasperry Pi 3B. The fish are video-recorded from the ventral side.

(B) Optomotor response (OMR) of medaka under the 940 nm LED light (red dots) and visible light (blue and gray dots). The direction of cylinder rotation is defined as the positive direction, and round number indicates the number of times the medaka swam in (+) or against (-) the direction of cylinder rotation. Since none of the medaka exhibited OMR throughout the dark period of the observation time (6:30 AM to 7:50 AM), it follows that the medaka do not perform sexual behavior in the dark (see Fig. 3) dependent on the dark-adapted vision.

Supplementary document S1. PMSG treatment and analysis

We performed intraperitoneal injection of pregnant mare serum gonadotropin (PMSG) (ASKA Animal Health Co. Ltd., Tokyo, Japan) to *gnrh1*$^{-/-}$ females for inducing ovulation according to Takahashi *et al.*,[17]. *gnrh1*$^{-/-}$ females were anesthetized with 0.02% Ethyl 3-aminobenzoate methanesulfonate salt (MS222) (Sigma-Aldrich, St. Louis, MO) and were injected with 10 U (10 μL) PMSG into their abdominal cavity using a syringe (Hamilton, Nevada, USA). We also injected the same volume of phosphate buffered saline (PBS; Takara Bio, Shiga, Japan) into the other *gnrh1*$^{-/-}$ females for negative control. The injected females were paired with WT male, and we recorded their sexual behavior by the same method for recording sexual behavior of KO female (Figure 1). When they spawned, the eggs were collected, placed to the breeding water containing methylene blue (JAPAN PET DESIGN, Tokyo, Japan). We calculated the fertilization rate by dividing the number of fertilized eggs by the total number of collected eggs.

Supplementary document S2. Analysis of optomotor response under light/dark conditions

To examine whether 940 nm wavelength light is invisible for medaka, we performed the optomotor response experiments. We prepared 18 cm-diameter circular tank with 3 cm water depth, placed medaka therein, and placed a transparent cylinder at the center of the tank to make the object fish swim in the periphery of the tank. The tank was enclosed by a paper cylinder with black and white stripe patterns of 2 cm width. The paper cylinder can be rotated around the tank at an arbitrary time, which was regulated by the DC motor (RF300CV-11320-23.5M-R, Shenzhen, China) attached to the Raspberry Pi 3 Model B equipped with Raspberry PiNoir Camera Module V2.1 via relay unit. The LED light of which wavelength is 940 nm (FRS5JS) was placed above the tank, which was turned on during the dark period (22:00 PM-8:00 AM).

At 6-7 PM of the previous day of behavioral testing, the WT medaka (body weight (g): 0.177～0.302) was placed into each tank, and were recorded for 10 seconds at 6:30/7:30/7:50 AM (dark periods) and 8:30 AM (light period) with the paper cylinder rotating. We counted the number of times the object fish swam around the tank.

Supplementary Video 1. Representative movie showing the sexual behavior exhibited during the light period (Related to Figure 3)

From the start of the video (0:05), the male (below) approached the female (above) from behind and followed her at the same speed (following, 0:05~), and subsequently the male exhibited circular turn in front of the female (courtship display, 0:08~). After the several times of following and courtship display, the male and female contacted with each other and quivered together (clasping, 1:43 ~ 2:12). Spawned eggs can be observed on the female abdomen (2:20).

Supplementary Video 2. Representative movie showing the sexual behavior exhibited during the dark period (Related to Figure 3)

From the start of the video (0:05), the male (bottom) and the female (surface) swam in different directions, keeping their distance from each other. The male approached the female from behind and followed her at the same speed (following, 0:46 ~), and the male and female contacted with each other and quivered together (clasping, 1:15 ~ 1:32). We can observe the spawned eggs attached to the female abdomen (2:45).
